# ConsensusDriver improves upon individual algorithms for predicting driver alterations in different cancer types and individual patients – a toolbox for precision oncology

**DOI:** 10.1101/127985

**Authors:** Denis Bertrand, Sibyl Drissler, Burton Chia, Jia Yu Koh, Li Chenhao, Chayaporn Suphavilai, Iain Beehuat Tan, Niranjan Nagarajan

## Abstract

**Background:** In recent years, several large-scale cancer genomics studies have helped generate detailed molecular profiling datasets for many cancer types and thousands of patients. These datasets provide a unique resource for studying cancer driver prediction methods and their utility for precision oncology, both to predict driver genetic alterations in patient subgroups (e.g. defined by histology or clinical phenotype) or even individual patients.

**Methods:** We performed the most comprehensive assessment to date of 18 driver gene prediction methods, on more than 3,400 tumour samples, from 15 cancer types, to determine their suitability in guiding precision medicine efforts. These methods have diverse approaches, which can be classified into five categories: <u>f</u>unctional <u>i</u>mpact on proteins in general (FI) or specific to <u>c</u>ancer (FIC), <u>c</u>ohort-<u>b</u>ased <u>a</u>nalysis for recurrent mutations (CBA), <u>m</u>utations with <u>e</u>xpression <u>c</u>orrelation (MEC) and methods that use gene <u>i</u>nteraction <u>n</u>etwork-based <u>a</u>nalysis (INA).

**Results:** The performance of driver prediction methods varies considerably, with concordance with a gold-standard varying from 9% to 68%. FI methods show relatively poor performance (concordance <22%) while CBA methods provide conservative results, but require large sample sizes for high sensitivity. INA methods, through the integration of genomic and transcriptomic data, and FIC methods, by training cancer-specific models, provide the best trade-off between sensitivity and specificity. As the methods were found to predict different subsets of drivers, we propose a novel consensus-based approach, ConsensusDriver, which significantly improves the quality of predictions (20% increase in sensitivity). This tool can be applied to predict driver alterations in patient subgroups (e.g. defined by histology or clinical phenotype) or even individual patients.

**Conclusion:** Existing cancer driver prediction methods are based on very different assumptions and each of them can only detect a particular subset of driver events. Consensus-based methods, like ConsensusDriver, are thus a promising approach to harness the strengths of different driver prediction paradigms.

## Background

Cancers result from the accumulation of various types of DNA mutations including point mutations, indels, large scale copy number aberrations (CNA), and structural variations [1]. During tumor development, in addition to mutations that confer functional advantages to tumor cells (i.e. driver mutations) [2], a large number of passenger mutations with no or little functional impact may arise, confounding our ability to identify the key events in oncogenesis for understanding and treating cancers [3].

Recent large scale cancer genome sequencing efforts such as The Cancer Genome Atlas (TCGA) [4], International Cancer Genome Consortium (ICGC) [5] and other studies (e.g. [6, 7]) have harnessed technological advances in DNA/RNA sequencing to provide comprehensive mutation catalogs and associated omics profiles in tumors. These compendiums provide a rich resource for the development of integrative cancer driver prediction methods [8–10]. In addition, they further highlight the challenges that still remain in driver prediction. In particular, due to the heterogeneity of cancer types, often few frequently mutated (and likely driver) genes were identified in these studies with many more genes being rarely mutated and thus indistinguishable from noise due to passenger mutations [11–13]. Despite this, the ability to identify cancer drivers may be key for improved targeted therapy [14–16]. For example, breast cancer patients with ERBB2 driver mutations can respond successfully to the ERBB2 inhibitor trastuzumab [17], but similar therapy may also benefit patients with other cancers where ERBB2 mutations are rare [18]. After the initial wave of large-scale cancer studies, different cohorts of patients continue to be sequenced with more distinct phenotypes (e.g. previously unprofiled disease sites or disease states such as tumors characterized by primary or acquired drug resistance). In the growing paradigm of precision oncology individual patients are also sequenced either broadly (whole exome) or with targeted sequencing panels of genes selected by above studies and existing knowledge about the patient’s disease and available treatments, in order to gain insights into biology and to match the right patient to the right drug at the right time. There is thus a deep biological and clinical need to identify the mutation that drives the tumor of a single patient.

Due to its biological and practical importance, a range of different approaches have been proposed for inferring the impact of mutations on genes and their likely role in cancer. These methods differ widely in the information they require as input (e.g. point mutations, indels, CNAs, expression data etc.) and in the models/assumptions that they use [19–21]. For example, many methods are based on using information about protein structure and evolution to detect point mutations that may have a <u>f</u>unctional <u>i</u>mpact in general (FI) [22–25], or specifically in the context of <u>c</u>ancer (FIC) [26–28]. These methods predict functional/driver mutations in each sample independently and their relative strengths have been studied in previous work [29, 30]. With the availability of large and heterogeneous cancer genomic datasets, newer methods have focused on <u>c</u>ohort <u>b</u>ased <u>a</u>nalysis to search for biases in mutation frequency indicative of positive selection in driver genes (CBA) [31–36] (compared in [37]), or mined for <u>m</u>utation-<u>e</u>xpression <u>c</u>orrelations to highlight driver CNAs (MEC) [38–40] (jointly evaluated in [41]). Other approaches have used mutual exclusivity of driver mutations to identify them in a large cohort of patients [42, 43]. Finally, a few methods have sought to incorporate information about gene <u>i</u>nteraction <u>n</u>etworks in their <u>a</u>nalysis with the aim of providing more sensitive predictions [44, 45], or to enable driver prediction based on integrative analysis of genomic and transcriptomic data (INA) [8–10].

Despite the diversity of driver prediction methods, a comprehensive evaluation of the strengths and weaknesses of different classes of methods on a diverse range of cancer types has not been conducted. We sought to address this by evaluating the performance of a panel of 18 different computational methods, covering a wide variety of models and input data types, on >3,400 tumor datasets from 15 TCGA cancer types. Methods were evaluated systematically for their concordance with gold-standard lists of driver and passenger genes, for their robustness to noise in the input, for their utility for working with data from small patient cohorts and for their ability to provide accurate and actionable patient-specific predictions for precision medicine applications. The overall predictive power for driver events was found to be moderate, highlighting the need for novel approaches and improved methods. Additionally, predictions from different classes of methods were found to be orthogonal to each other, motivating the development of a consensus-based approach (ConsensusDriver) to increase sensitivity and specificity of driver predictions across cancer types. Consensus-based approaches such as ConsensusDriver provide a systematic way to combine the strengths of different driver prediction algorithms in building an analytical toolbox for precision oncology.

## Results

### Different cancer types represent diverse driver prediction challenges

For the purpose of this study, we selected 15 different cancer types from TCGA for which exome sequencing, copy number and expression data (RNA-seq or arrays) were available (BLCA: Bladder Urothelial Carcinoma, BRCA: Breast Invasive Carcinoma, COAD: Colon Adenocarcinoma, GBM: Glioblastoma Multiforme, KIRC: Kidney Renal Clear Cell Carcinoma, KIRP: Kidney Renal Papillary Cell Carcinoma, LIHC: Liver Hepatocellular Carcinoma, LUAD: Lung Adenocarcinoma, LUSC: Lung Squamous Cell Carcinoma, OV: Ovarian Serous Cystadenocarcinoma, PAAD: Pancreatic Adenocarcinoma, PRAD: Prostate Adenocarcinoma, READ: Rectum Adenocarcinoma, STAD: Stomach Adenocarcinoma, THCA: Thyroid Carcinoma; see **Methods**). The cancer types selected vary widely in cohort sizes, mutational burden per patient and distribution of mutation types, thus representing a diverse set of challenges for driver prediction methods (**Figure 1a**). For example, we noted that while some cancer types are predominantly affected by point mutations (KIRP) or CNAs (OV), others have similar number of genes affected by both point mutations and CNAs (GBM). In addition, certain cancer types exhibited a bimodal distribution for mutational burden (READ, COAD, PRAD and KIRC) and this could impact the distributional assumptions of some methods. The distribution of mutation frequencies across genes also showed high variation between cancer types (**Supplementary Figure 1**). For example, while LUSC and OV have many genes with mutation frequency above 25%, THCA has only 3 genes with frequency above 5%, potentially impacting the sensitivity of methods that are dependent on mutation frequency for driver prediction (**Figure 1b**). We additionally noted that most tumors exhibited both point mutations and CNAs (**Supplementary Figure 2a**) and thus methods that take only a subset of mutation types as input may be at a disadvantage in terms of sensitivity (e.g. FI and FIC methods which only consider missense variants; **Supplementary Figure 2b**).

**Figure 1:**
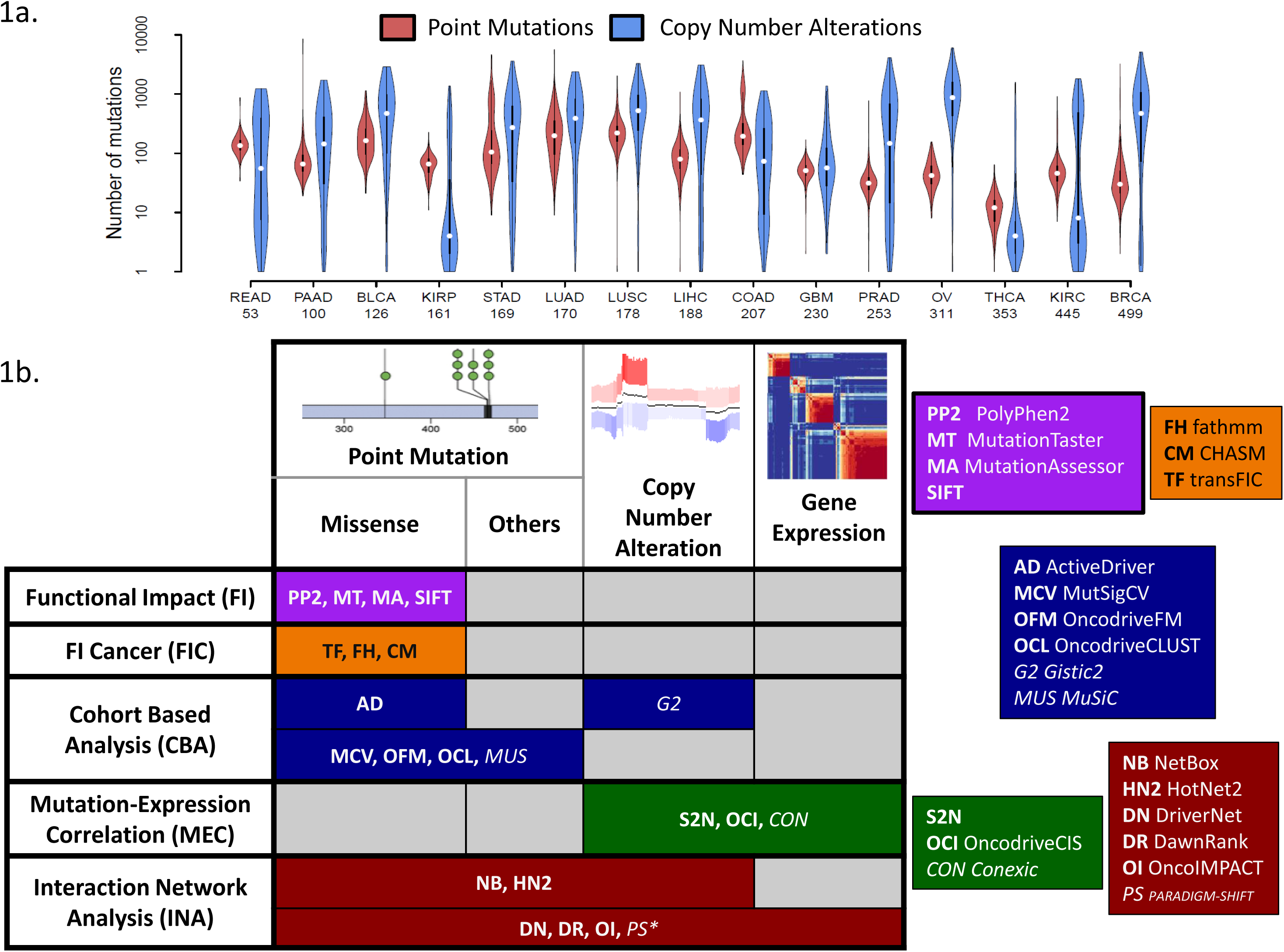
Diversity of driver prediction methods and datasets. (a) Violin plots showing that cancer types vary widely in terms of their point mutation and CNA burden (number of patient is indicated under the name the cancer types). (b) Two-way classification of driver prediction methods based on input data-types and modeling assumptions/approaches. The italicized methods were not included due to practical constraints on running them. * - PARADIGM-SHIFT is restricted to the analysis of point mutations.

In terms of driver prediction methods, we attempted to be as comprehensive as possible, though some methods had to be excluded due to feasibility issues (see **Methods**). In total, we evaluated 18 methods that could be used for driver prediction (**Figure 1b**), classifying these methods into (i) methods that belong to the Functional Impact (FI) category (primarily designed to identify function altering mutations but have been used for predicting drivers [29, 30, 46]) such as SIFT [22], PolyPhen2 (PP2) [23], MutationTaster (MT) [24] and MutationAssessor (MA) [25], (ii) methods that tailor this idea to cancer by learning specific models (Functional Impact in Cancer; FIC) such as CHASM [26], transFIC (TF) [27] and fathmm (FH) [28], (iii) methods that use cohort based analysis to detect signals of positive selection (CBA) such as ActiveDriver (AD) [36], MutSigCV (MCV) [31], MuSiC (MUS) [32], OncodriveCLUST (OCL) [33] and OncodriveFM (OF) [34] (all point mutation based), (iv) methods that integrate mutation data with transcriptomic data by looking for mutation-expression correlations (MEC) such as Conexic (CON) [38], OncodriveCIS (OCI) [39] and S2N [40], and finally (v) methods that use information from gene/protein interaction networks to analyze the effect of mutations such as NetBox (NB) [44], HotNet2 (HN2) [45], DriverNet (DN) [8], DawnRank (DR) [9] and OncoIMPACT (OI) [10]. We evaluated these 18 methods in predicting cancer drivers in patient cohorts and in individual patients.

### Driver gene prediction identifies many novel drivers but sensitivity is still a bottleneck

To evaluate the ability of various driver prediction methods to accurately differentiate between driver and passenger genes in a dataset, we compiled gold-standard lists for both. Specifically, we took the union of 5 different curated lists of drivers that have been reported before, including the widely used Cancer Gene Census list [47], a manually curated list of driver genes affected by copy number alterations [48], genes annotated as oncogenes by UniProt [49], a gene list derived from the Vogelstein 20/20 rule [11], and a gene list derived from literature mining [50] (**Supplementary Table 1**). Passenger genes were defined by taking the union of two manually curated lists of known passengers from NCG4 [51] and Rubio-Perez et al. [52] (**Supplementary Table 1**). These gold-standards are limited in that they are not cancer type or sample specific (though drivers are frequently shared [31, 53] and targeted [54] across cancer types), but represent an attempt to construct as comprehensive a list as possible such that novel cancer driver genes can be more effectively demarcated. The methods were evaluated on how well their predictions identified cancer driver genes based on three standard measures (as well as others as detailed below): precision (fraction of predictions that belong to the gold standard), recall (fraction of the gold standard contained in the predictions) and the F1 score that combines both precision and recall (see **Methods** for a more detailed description).

Due to the wide variation in the number of driver predictions from different methods (median of 10 for MutSigCV to >8,000 for MutationTaster; **Supplementary Figure 3**) we restricted our analysis to either the top 10 or top 50 predictions from each method (see **Supplementary Note 1** and **Supplementary Figure 4** for further details). An overview of the top 50 predictions for each method can be seen in **Figure 2a**. In general, most methods report a low number of passenger genes in their top 50 predictions except for FI methods (~20% of predictions). This is as expected as FI methods are not designed to specifically exclude function altering mutations that may not be linked to cancer, unlike the FIC methods.

**Figure 2:**
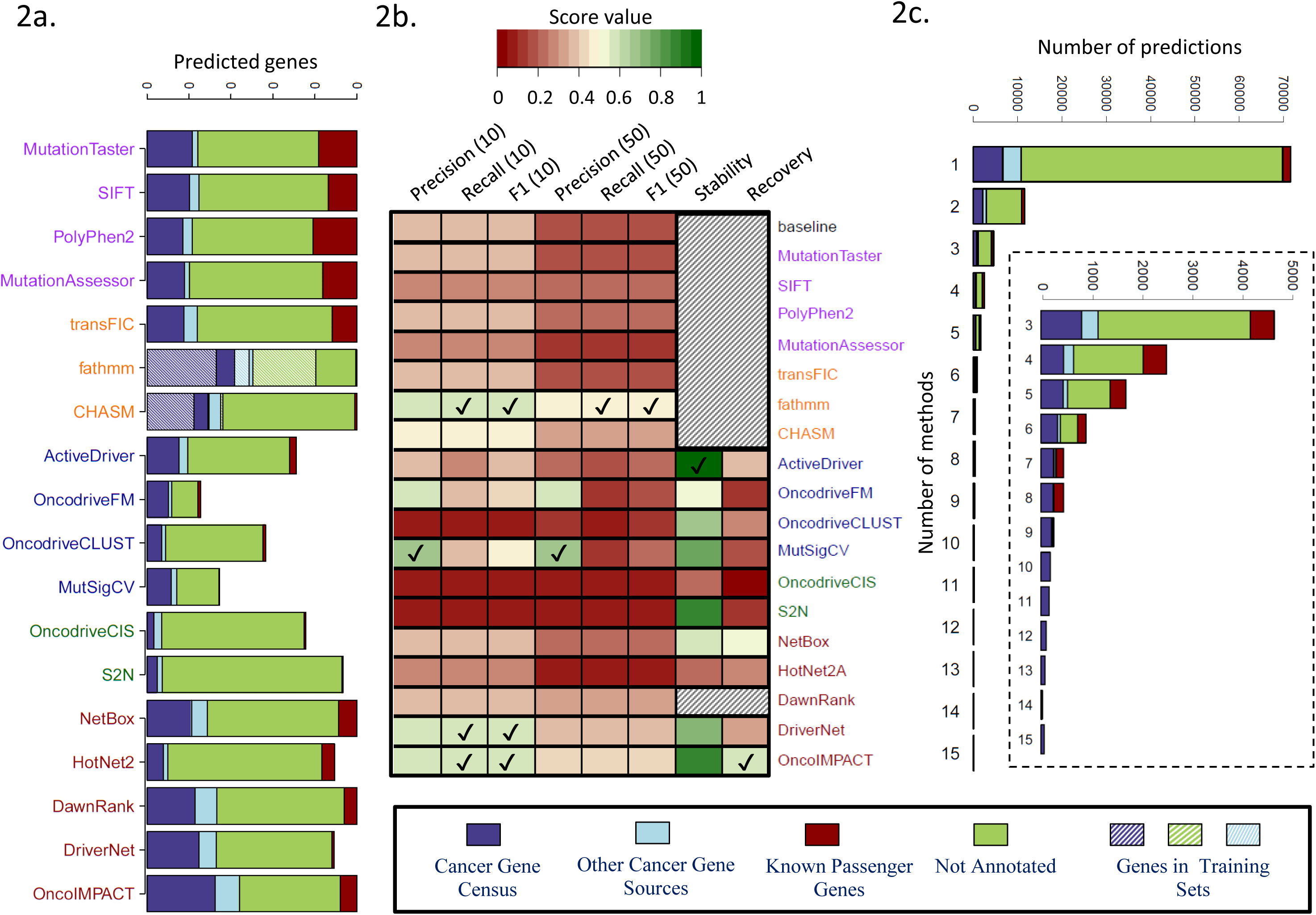
Evaluation of cohort-level predictions across cancer types. (a) Average number of genes (over 15 cancer types) among top 50 predictions that belong to different classes (known drivers, passengers and other genes). Note that some methods have less than 50 predictions on average. (b) Summary results for the evaluation of the 18 driver prediction methods according to various criteria: precision (for top 10 or top 50 predictions), recall (for top 10 or top 50 predictions) and F1 score based on comparison with gold-standard of known cancer genes, stability (precision when evaluated on predictions from the full data set) and recovery (recall when evaluated on the full data set) based on down-sampling to a dataset of 50 samples. Results are averaged over 3 replicates on the 7 cancer types with ≥200 samples. For each evaluation metric, methods with the highest score are indicated by a tick mark. Shaded cells represent methods for which the down-sampling was not performed, either because they are not affected by down-sampling (baseline, FI and FIC methods) or due to high computational time requirements (DawnRank). (c) Annotation of the predicted driver genes according to the number of methods they are predicted by.

The number of known cancer-associated genes reported in the top 50 predictions of different methods varied widely, from a mean of 4 for OncodriveCIS to 27 for fathmm (a majority of these belong to the Cancer Gene Census). In general, the highest sensitivity was provided by methods in the FIC and INA categories, reporting >15 known drivers in the top 50 list. Note that the FIC methods use a machine learning approach with training sets that substantially intersect our gold standard (**Figure 2a**), and thus their sensitivity to predict new drivers may not be accurately captured here. On the other hand, methods in the CBA category were most concordant with the list of gold standard drivers (0.5 and 0.6 for OncodriveFM and MutSigCV respectively; **Figure 2b** and **Supplementary Figure 5**). Selecting the best method in each category, we observed that all methods were more enriched for drivers in their top predictions as expected, and methods such as fathmm and OncoIMPACT retained high precision even for predictions lower down the list (**Supplementary Figure 6**). A striking aspect of the results in **Figure 2a** is the large number of predicted genes that are neither passengers nor known driver genes. The majority of these genes are predicted by a single method and are likely enriched in false positives (**Figure 2c**). However, genes predicted by multiple methods were strongly enriched in cancer related functions (**Supplementary Figure 7b**), highlighting the fact that many more driver genes remain to be discovered, and consistent with recent work showing that more driver genes exist even in extensively studied TCGA cancer types [13].

We used the F1 score that combine precision and recall to rank methods and compare them against a “baseline” method that simply orders genes based on mutation frequency (**Figure 2b**, **Supplementary Figure 8**; see Methods). The methods fathmm, CHASM, NetBox, DawnRank, DriverNet and OncoIMPACT provided significantly better results than baseline for precision and F1 score, while ActiveDriver, OncodriveFM and MutSigCV showed significant improvement in precision (Wilcoxon rank sum test *p*-value < 0.1; **Supplementary Figure 9**). The lower scores observed for MEC methods was not explained solely by their restriction to CNAs (**Supplementary Figure 10**).

To evaluate how the driver predictions are affected by cohort size, we tested the different methods for robustness and power using a subsampling approach that compares predictions for a method to those on the full dataset (stability = precision and recovery = recall compared to results from full dataset; see Methods) [10]. Many methods exhibited high stability (>70%) at least for the 50 and 100 sample comparisons (ActiveDriver, MutSigCV, S2N, DriverNet, OncoIMPACT; **Figure 2b** and **Supplementary Figure 11a**). However, few methods exhibited recovery >50% (NetBox, OncoIMPACT), highlighting challenges in uncovering drivers when cohort sizes are limited (**Figure 2b** and **Supplementary Figure 11b**). Overall, as summarized in **Figure 2b**, no single method outperformed the others in all metrics.

### Most methods predict no drivers for 10% of patients but many provide robust patient-specific predictions

We next evaluated methods for their ability to accurately identify driver events in a patient-specific manner to assess their utility for precision medicine applications. Note that not all methods provide predictions per patient and for such methods, we assumed that nominated driver genes are drivers in all patients in which they were mutated. We began by computing statistics for the number of drivers nominated in each patient by various methods, under the assumption that reporting too few (<1) or too many drivers (>15) may make them less useful (**Figure 3a**; number of drivers per patient is generally expected to be <10 [1, 55, 56]). Interestingly, with the exception of FI methods that call a large number of drivers for the majority of the patients, nearly all the other methods report no driver events for >10% of patients. This could be an indication of low sensitivity but could also be due to driver events having other origins (e.g. copy-number neutral rearrangements, large translocations, regulatory or noncoding mutations or methylation and other epigenetic events) that were not considered by these methods. The method OncoIMPACT was found to be unusual in this aspect (even compared to INA methods) as it identified at least 1 driver in nearly all patients. Methods belonging to the CBA and MEC categories typically identified <2 drivers in a large fraction of the cohorts (~40%). On the other end, some methods frequently (in >50% of cohort) identified >15 drivers in patients, suggesting that they may be overcalling at the patient-specific level (MutationTaster, MutationAssessor, SIFT, PolyPhen2 and S2N; **Figure 3a**).

**Figure 3:**
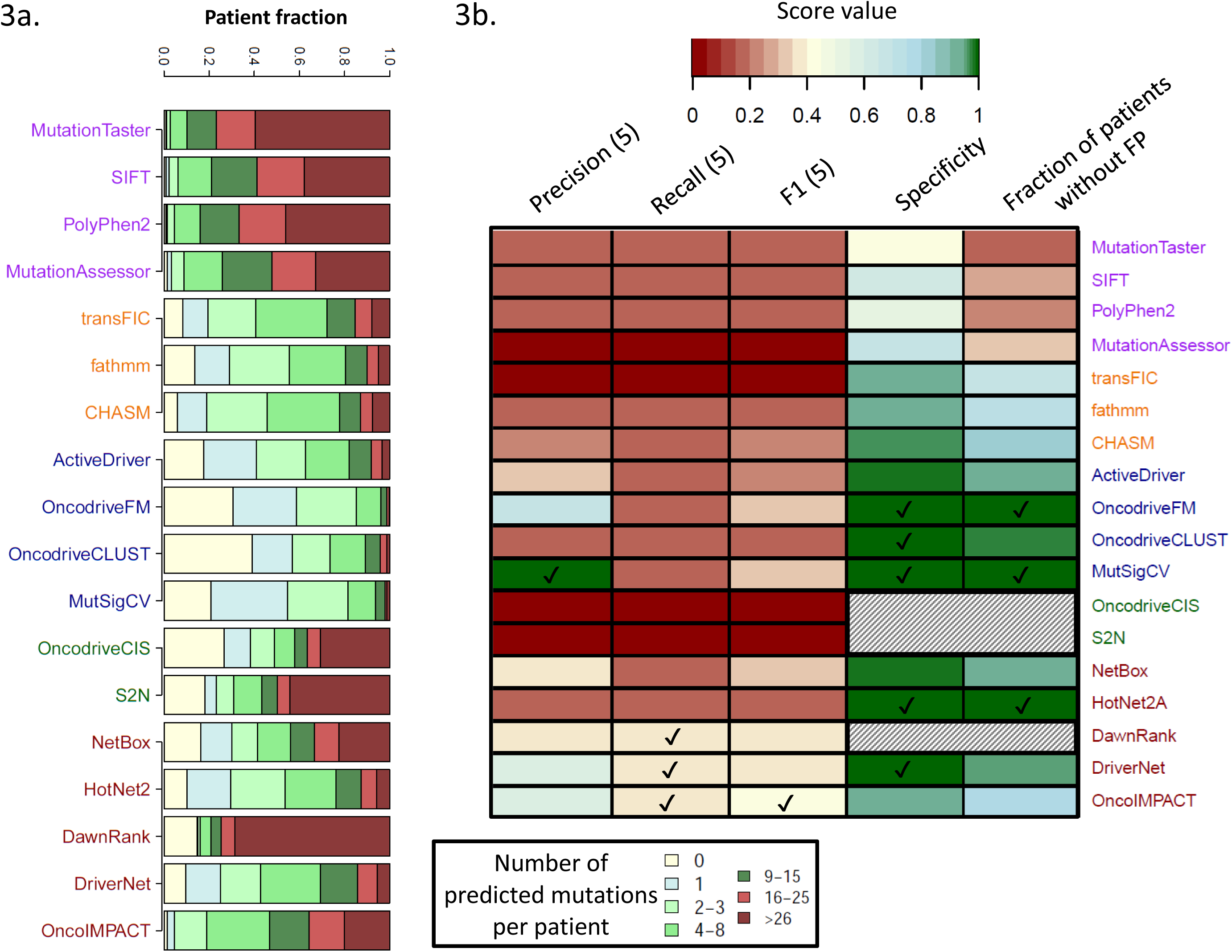
Evaluation of patient-specific predictions. (a) Number of predicted drivers per patient. DawnRank was excluded for this analysis as it reports all mutations for a patient with no filtering criteria provided. (b) Summary results for the evaluation of the 18 driver prediction methods according to various criteria: precision, recall and F1 score based on comparison with gold-standard of known cancer genes (for top 5 patient-specific predictions), specificity and fraction of patients without false positives (FP) in their top 5 predictions based on the introduction of decoy mutations (10% of the number of mutations in the tumor). Results are averaged over 3 replicates for each cancer type. For each evaluation metric, methods with the highest score are indicated by a tick mark. Shaded cells represent methods for which the introduction of missense decoy mutations was not performed, either because they only process CNA data (S2N and oncodriveCIS) or due to high computational time requirements (DawnRank).

Considering the top 5 patient-specific predictions, most methods provided similar precision and F1 score as in the cohort-level evaluation, with the network based methods (INA) generally outperforming other approaches (**Figure 3b** and **Supplementary Figure 12**). As before, CBA methods such as OncodriveFM and MutSigCV provided the best precision (**Figure 3b** and **Supplementary Figure 13**).

To test robustness to noise and to estimate the specificity of the predictions at the patient-specific level, we introduced decoy passenger mutations in genes with probability weighted by the gene length (see **Methods**). Most methods exhibited good robustness to such noise with specificity generally higher than 95% (except for FI methods; **Figure 3b** and **Supplementary Figure 14a**). In particular, methods in the FIC category accounted well in identifying decoy function altering mutations, improving significantly over methods in the FI category. Also, since CBA methods explicitly model such sources of noise, they were found to have the best control over them. We also noted that most patients (>80%) do not have any of the decoy mutations in their predictions even when the overall specificity of a method is ~95% (**Figure 3b** and **Supplementary Figure 14b**). This is even more the case when only the top 5 or 10 predictions are considered, highlighting the robustness of many methods at the patient specific level. As summarized in **Figure 3b**, no single method uniformly outperformed the others at the patient-specific level as well.

### Prioritization of actionable drivers is still a challenge for most individual methods

The prioritization of driver genes and mutations that are actionable is a key requirement for decision support systems to aid in precision oncology. We sought to evaluate the performance of the various methods studied here based on curated lists of actionable genes (genes that can be targeted by a drug under certain conditions) from the OncoKB [57] and IntOGen [52] databases (see **Methods** and **Supplementary Table 2**). Analyzing the top 5 driver predictions per patient from each method, we observed significant variability in performance, with the fraction of patients with a predicted actionable driver varying from 6% for OncodriveCIS to >60% for DriverNET and OncoIMPACT (**Figure 4a**). We observed that the different methods provided largely non-overlapping predictions, enabling the union to predict actionable drivers for up to 81% of patients. A breakdown of the predictions by cancer type (**Figure 4b**) highlighted that six of them (LIHC, PRAD, KIRP, OV, KIRC, BRCA) have a much lower fraction of patients with predicted actionable driver genes. This could in part be due to the lack of sensitivity in driver prediction methods, but in most cases it is explained by the cancer types being enriched for non-targetable drivers, highlighting the need for further drug discovery efforts in these cancer types. Finally, as a positive control test, we assessed the sensitivity of the methods in predicting two known actionable oncogenes, BRAF (various drugs are FDA approved for treating melanoma with V600 mutations [58]) and PIK3CA (the inhibitor alpelisib is currently undergoing a clinical trial for breast cancer [59]) in patients harboring known oncogenic mutations (BRAF V600 and 19 PIK3CA mutations located in the domains of the catalytic subunit [57], see **Supplementary file 3**). We observed notable variation in the numbers of patients where the mutations were flagged as drivers by different methods (**Figure 4c**), with multiple methods that did not report the genes for any patient (similar results were observed with top 10 predictions; **Supplementary Figure 15**). These results highlight that the differences in the underlying model of various methods can lead to dramatically different abilities in predicting actionable driver genes and that care should be exercised in interpreting and integrating results from different driver prediction systems.

**Figure 4:**
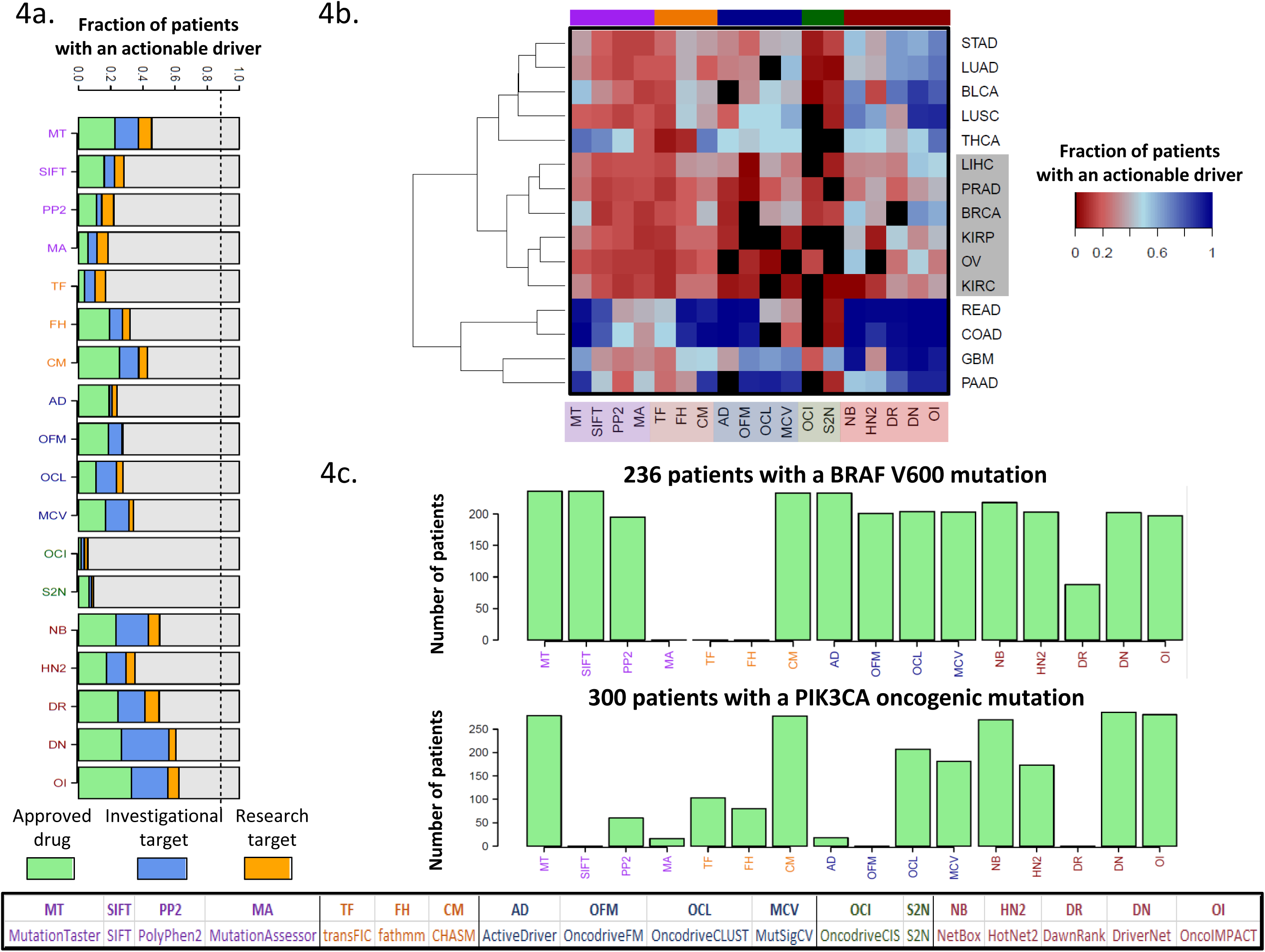
Sensitivity of methods for identifying actionable drivers. Results are reported based on the top 5 driver predictions for each method. (a) Fraction of patients with at least one actionable driver gene predicted. The dashed line represents the fraction of patient with at least one mutated actionable gene. The dashed line represents the fraction of patients where an actionable gene is mutated. (b) Breakdown of patients according to their cancer type reveals a cluster of 6 cancer types (highlighted in grey rectangle) with a low fraction of patients with predicted actionable drivers (complete linkage hierarchical clustering using Euclidean distance) (c) Number of patients with a predicted actionable driver mutation for the genes BRAF and PIK3CA.

### Low concordance across methods enables the construction of a better consensus-based approach

A comparison of driver gene predictions across methods revealed that in addition to the expected differences across categories, many methods had a significant number of calls that were unique to them (**Supplementary Figure 16a**). This was particularly the case for FIC methods such as fathmm and CHASM, and network based methods (INA) such as DawnRank, DriverNet and OncoIMPACT. In addition, for the more sensitive methods (e.g. fathmm and OncoIMPACT), many predictions were shared by >4 methods suggesting that this could provide additional confidence for many of their calls. To evaluate if consensus approaches could improve over predictions from individual methods, we evaluated a rank-aggregation based approach [60] using all methods (BORDAall) as well as a subset of methods identified using cross-validation (ConsensusDriver; see **Methods**). We found that the same methods were consistently selected by ConsensusDriver across cancer types, covering a wide range of methods across categories (**Figure 5b**, **Supplementary Figure 17**), including CHASM, fathmm (FIC), OncodriveFM, MutSigCV (CBA), DriverNet and OncoIMPACT (INA).

**Figure 5:**
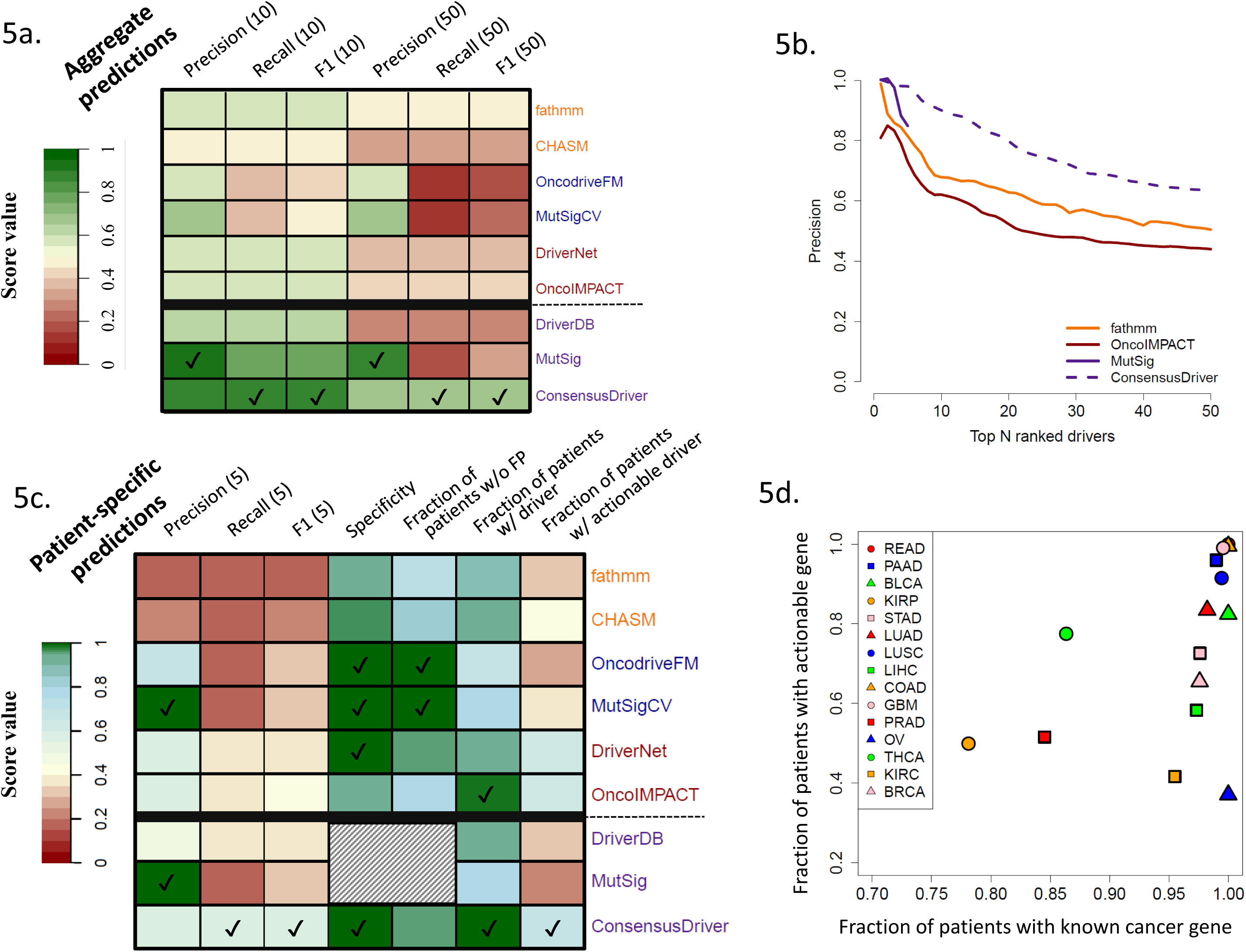
The utility of a consensus approach for driver prediction. (a) Comparison with gold standard of known cancer genes (precision, recall and F1 for the top 10 and 50 prediction) of the cohort-level predictions of ConsensusDriver, the 6 methods that it integrates (CHASM, fathmm, OncodriveFM, MutSigCV, DriverNet and OncoIMPACT), and consensus gene lists from DriverDB and MutSig. (b) Precision (fraction of predictions that belong to the gold standard) as a function of the number of predictions. (c) Evaluation of patient-specific prediction for methods presented in **Figure 5a** according to various criteria: precision, recall and F1 score based on comparison with gold-standard of known cancer genes (for top 5 predictions), specificity and fraction of patients without false positives (FP) in their top 5 predictions based on the introduction of decoy mutations (10% of the number of mutations in the tumor) and the fraction of patients with at least one predicted driver or actionable gene in the top 5 predictions. For each evaluation metric, methods with the highest score are indicated by a tick mark. Shaded cells represent experiments that we were not able to perform on DriverDB and MutSig as they only provide gene lists. (d) Scatter plot depicting the number of known cancer genes and actionable genes in the top 5 patient-specific predictions of ConsensusDriver across cancer types.

Across cancer types, while ConsensusDriver was able to improve over the best individual methods (1.4× improvement compared to fathmm in median F1 score, one-sided Wilcoxon rank sum test *p*-value = 10^−3^; **Figure 5b** and **Supplementary Figure 16b**), BORDAall did not show a significant improvement in precision (**Supplementary Figure 18**) or in the F1 score (**Supplementary Figure 16b**; see **Supplementary Note 2** and **Supplementary Figure 19** for comparisons with other machine learning approaches). Comparing ConsensusDriver to two consensus-based gene lists, we noted that it improved recall and F1 performance over both of them (MutSig [13] and DriverDB [61]; one-sided Wilcoxon rank sum test *p*-value = 10^−3^ and ^2×10−4^ for F1 improvement). Overall ConsensusDriver is a consistent improvement over individual methods and consensus-based gene lists exhibiting a precision of 0.9 for its top 10 predictions and 0.63 over its top 50 predictions (**Figure 5b**).

At the sample-specific level, ConsensusDriver is largely better than individual methods across metrics (e.g. it provides 1.5× improvement over OncoIMPACT in precision [one-sided Wilcoxon rank sum test *p*-value < 2×10^−16^] and 1.35× improvement in F1 score [one-sided Wilcoxon rank sum test *p*-value = 2.4× 10^−16^]; **Figure 5c**), with the exception of precision (versus MutSigCV and OncodriveFM) and the fraction of patients without false positive predictions (versus MutSigCV). It arguably provides a better tradeoff though, by improving the fraction of samples with a predicted driver gene (from 0.8 for MutSig to 0.99) and predicted actionable driver genes (from 0.36 for MutSigCV to 0.67). This improved sensitivity is also accompanied by high specificity (0.99) for ConsensusDriver (**Figure 5c**). The additional sensitivity of ConsensusDriver helped establish that, with the exception of THCA, PRAD and KIRP, most of the patients analyzed here have at least a known cancer gene in their predictions (**Figure 5d**). The fraction of patients with actionable predicted driver genes is however lower, as many known driver genes are still not targetable (e.g. Ovarian Cancer, where most patients harbor a TP53 mutation, exhibits the lowest fraction of patients with a predicted actionable gene).

## Discussion

We provide the first systematic evaluation of different classes of driver prediction methods over a large number of cancer types. As the community still lacks standard evaluation protocols, we identified various criteria to evaluate predictions at the cancer-type level (concordance and sensitivity over know cancer genes, and stability/recovery of predictions upon sub-sampling) and at the patient-specific level (number of driver genes per patient, concordance with gold standard, robustness to noise mutations). The availability of our pre-formatted datasets, predictions from evaluated methods as well as a package of tools to study new predictions, provides a useful resource and a standardized framework to evaluate any newly developed method against a diverse panel of state-of-the-art methods and on a large number of cancer types.

A key result of our analysis is that there is no single method (or category of methods) that generally outperforms other methods and instead there are specific pros and cons that need to be taken into consideration when selecting a method for analyzing new datasets. For example, FIC methods are more appropriate for the analysis of a small number of samples when only exome data is available, while CBA methods should be selected for large-scale exome sequencing data sets and INA methods provide greater sensitivity when genomic and transcriptomic data is available. In general, our study highlights the value of integrative methods: for example, methods that are restricted to point mutations, not surprisingly, have a large drop in sensitivity in cancer types with significant amount of CNA events. In the ovarian cancer dataset, the best CBA method only predicts 3 known cancer genes compared to 18 using the best INA method. Furthermore, INA methods that integrate expression data (DriverNet, DawnRank and OncoIMPACT) show, in most analyses, better results than methods that analyze only genomic data (NetBox and HotNet2). Further work is thus needed in this area, particularly in developing methods that incorporate information from other data-types (e.g. miRNA-seq) and other mutation types (e.g. non-coding mutations).

Our study also provides a detailed analysis of the driver predictions at the patient level. It highlights the robustness (low false positive rate, high concordance with the gold standard) of the driver predictions, but also the lack of sensitivity (significant fraction of patients with 0 to 1 driver predicted) of the vast majority of methods, with methods integrating expression having the best performance. In terms of prioritizing actionable genes, most methods have even more severe limitations and integrating methods with different underlying models could help ameliorate this problem.

There are several limitations to our work: Firstly, we limited our analysis to a single data source (TCGA, which currently provides the most comprehensive coverage of cancer types with genomic and transcriptomic data) and to a set of well-cited methods with software implementations that we were able to use successfully. Secondly, our evaluations were based on gold-standard lists of driver genes that are not cancer-type or patient specific. They thus do not necessarily reflect the heterogeneity of cancers and lack direct evidence that a specific mutation has a functional role in a particular tumor. Other large-scale initiatives have tried to bridge this gap and provide cell-line specific shRNA (Achilles[62]) and drug resistance profiles (CCLE [63] and GDSC [64]). The results of these studies could potentially be used to generate more refined gold-standards. However, such analysis will not come without drawbacks as (i) the cell lines used typically do not have normal controls and thus mutation calls can be error-prone and (ii) the experiments are limited to measuring cell growth and thus miss other relevant phenotypes (e.g. motility, invasiveness etc.). Nevertheless, large experimentally derived and patient specific gold-standard driver gene lists are needed to further advance the development and evaluation of new driver prediction methods.

Overall, our study highlights that while existing driver predictions methods can have limited sensitivity as a function of data-types and modeling assumptions used, their diversity in fact provides an avenue for better consensus methods, as demonstrated by the novel consensus method proposed here (ConsensusDriver). Development of methods that harness new sources of information thus might provide greater benefits then refinement of existing paradigms for driver discovery. From a practical point of view, we provide an easy-to-use package to run 18 different driver prediction methods, as well as to aggregate their results into consensus predictions that are largely superior to the individual methods, thus serving as a valuable toolbox for precision oncology efforts.

## Material and Methods

### Data source and preprocessing

CNA and exome point mutation data for all cancer types was obtained from GDAC via Firehose (https://gdac.broadinstitute.org). All point mutations excluding synonymous mutations (i.e. indels, missense, nonsense and splice site variants) and CNAs with a value of 2 (focal amplification) or −2 (focal deletion) were used for downstream analysis. Expression data for tumor and normal samples for all cancer types was downloaded from the TCGA website (level 3; https://tcga-data.nci.nih.gov). For a detailed description of expression data analysis, see **Supplementary Methods**.

### Assessment of driver prediction methods

For each method, we used default parameters or the set of recommended parameters provided in the method’s manual or corresponding publication. In cases where methods required a threshold for candidate driver selection (e.g. on the *p*-value or score for candidates), we used the value indicated in the method’s publication or manual (see **Supplementary Methods** for a detailed description of the parameters and threshold used).

For analysis of patient-specific predictions, for most methods, mutated genes in each patient (with mutation types matching the expected input for the method) were ordered according to their rank on the full dataset. For FI and FIC methods, and for OncoIMPACT, mutation/patient specific scores were used to order genes (best score in the case of multiple mutations; ties broken by average gene score).

DriverDB predictions were obtained from http://driverdb.tms.cmu.edu.tw/driverdbv2/cancer.php and are based on the output of the following methods: ActiveDriver [36], Netbox [44], OncodriveFM [34], MutSigCV [31], Dendrix [43], MDPFinder [65], Simon [66] and MEMo [67]. Genes predicted by 2 or more methods were selected and ranked using the order provided on the DriverDB website. MutSig predictions were obtained from Lawrence et al [13].

A few methods were excluded from this benchmark for the following reasons: (i) they could not be run without further data processing, complex pre-filtering steps or inclusion of additional data (Genome MuSiC [32], Conexic [38], Mutex [42] and MultiDendrix [43]), (ii) they had prohibitively high computational requirements (PARADIGM-SHIFT [68]) or (iii) provided incompatible predictions (Gistic2 [35] with region-level predictions).

### Performance evaluation

#### Comparison with gold standard

We assessed the performance of all methods against a gold standard list of cancer driver genes (union of Cancer Gene Census [47], a manually curated list of CNA drivers [48], oncogenes from UniProt [49], gene list from the Vogelstein 20/20 rule [11], and a gene list from literature mining [50]) based on three different measures (based on the *N* top predictions): precision (*P*),

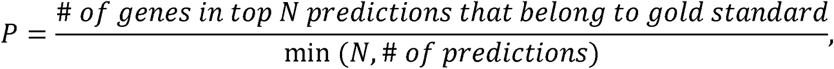
recall (R), 
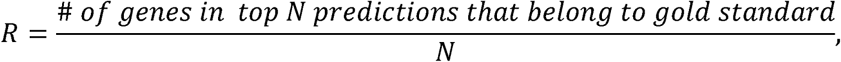
and the F1 score that combines both precision and recall, 
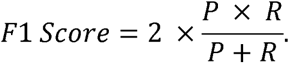

#### Robustness to subsampling

Subsampling analysis was performed for of each of the 7 cancer types with more than 200 tumor datasets. Two different measures were used to evaluate the robustness of results from a method on a subsample (*S*) when compared to its results on the full dataset (F): *stability* as a measure of precision when comparing the top *N* predictions of *S* (*S^N^*) to truth as defined by F,

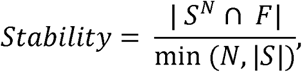
and *recovery* as a measure of sensitivity when comparing predictions in *S* to the top *N* predictions in *F* (*F^N^*),

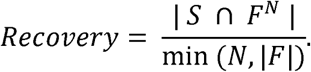

To make the comparison between and reasonable, we excluded from genes that were not mutated in the subsampled dataset. To avoid penalizing sensitive or conservative methods, we choose *N* to be 20 as a majority of the methods provided >20 predictions.

#### Generation of decoy missense mutations

For each patient, we introduced *n* false positive/decoy mutations, where *n* = 2%, 5% or 10% of the number of mutations in a tumor. Point mutations were randomly placed in coding regions of un-mutated genes with probability proportional to the coding length and missense mutations were selected using annovar (to avoid bias against methods that cannot analyze nonsense or splice-site mutations). For consistency, this analysis was restricted to the 12 cancer types annotated using the hg19 genome (i.e. COAD, OV and READ samples, annotated using hg18, were excluded).

#### Construction of an actionable gene list

We downloaded gene lists from IntOGen (https://www.intogen.org/) and OncoKB (http://oncokb.org/), and took the union of the actionable genes reported in them. We excluded drugs associated to a non-mutated gene from OncoKB, off-target genes from the IntOGen list, drugs targeting fusion genes, gene therapy targets and genes associated to drug resistance. Each drug/gene association was classified into three levels in the following order of preference: approved drug (Level 1 and 2A from OncoKB and “FDA approved drug” from IntOGen), investigational target (level 3A of OncoKB and “Drug in Clinical Trials” from IntOGen), and research target (all other genes).

### ConsensusDriver method

ConsensusDriver is based on the Borda approach, where each gene was given a score equal to the sum over all methods of either its rank, if the gene was ranked, or the maximum number of predictions in that data set (*M*), if the gene was unranked.

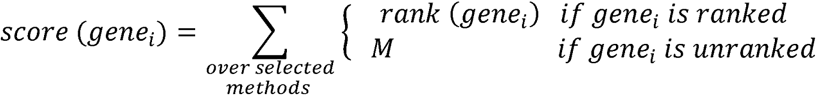

Genes were then re-ranked according to this score. To select the best set of methods for a particular cancer type, we used the following procedure (equivalent to a leave-one-out): (1) exhaustively compute the Borda consensus score on the 262,125 possible method combinations; (2) select the method combination which obtained the best average combine score for the others 14 cancer types. For the sample specific predictions, we integrated the patient specific-predictions of the six methods identified (fathmm, CHASM, OncodriveFM, MutSigCV, DriverNET and OncoIMPACT, **Supplementary Figure 17**). BORDAall used all the methods in constructing a BORDA based ranking.

## Availability of Supporting Data

A toolbox that contains scripts to reproduce results presented in this paper and evaluate results from newly developed methods is available at https://github.com/CSB5/driverevaluation. The site also contains results for each driver prediction method on all fifteen cancer types and the necessary input files (such as normalized expression, differential expression, mutations and copy number alteration lists). The ConsensusDriver package is freely available under the MIT license at https://hub.docker.com/r/csb5gis/consensusdriver and allows users to run individual driver prediction methods as well as the consensus algorithm.

## Competing Interests

The authors have no competing interests.

## Authors’ Contributions

DB, SD and NN designed the study. BC, DB, LC and SD implemented the scripts. DB, BC and SD conducted the experiments and analysis with guidance from NN. DB, BC, SC and NN drafted the manuscript with edits from IBT. All authors read and approved the final manuscript.

## Acknowledgements

This work was supported by funding from the Agency for Science, Technology and Research (A*STAR), Singapore. We thank Dr. Anders Skanderup and Dr. Asif Javed for insightful comments and suggestions on the manuscript.

## References

1. Stratton MR, Campbell PJ, Futreal PA: The cancer genome. Nature 2009, 458: 719–24.

2. Vogelstein B, Kinzler KW: Cancer genes and the pathways they control. Nat Med 2004, 10: 789–99.

3. Garraway LA, Lander ES: Lessons from the cancer genome. Cell 2013, 153: 17–37.

4. The Cancer Genome Atlas (TCGA) [http://cancergenome.nih.gov/]

5. Zhang J, Baran J, Cros A, Guberman JM, Haider S, Hsu J, Liang Y, Rivkin E, Wang J, Whitty B, Wong-Erasmus M, Yao L, Kasprzyk A: International Cancer Genome Consortium Data Portal‐‐a one-stop shop for cancer genomics data. Database (Oxford) 2011, 2011: bar026.

6. Shah SP, Roth A, Goya R, Oloumi A, Ha G, Zhao Y, Turashvili G, Ding J, Tse K, Haffari G, Bashashati A, Prentice LM, Khattra J, Burleigh A, Yap D, Bernard V, McPherson A, Shumansky K, Crisan A, Giuliany R, Heravi-Moussavi A, Rosner J, Lai D, Birol I, Varhol R, Tam A, Dhalla N, Zeng T, Ma K, Chan SK, et al.: The clonal and mutational evolution spectrum of primary triple-negative breast cancers. Nature 2012.

7. Zang ZJ, Cutcutache I, Poon SL, Zhang SL, McPherson JR, Tao J, Rajasegaran V, Heng HL, Deng N, Gan A, Lim KH, Ong CK, Huang D, Chin SY, Tan IB, Ng CCY, Yu W, Wu Y, Lee M, Wu J, Poh D, Wan WK, Rha SY, So J, Salto-Tellez M, Yeoh KG, Wong WK, Zhu Y-J, Futreal PA, Pang B, et al.: Exome sequencing of gastric adenocarcinoma identifies recurrent somatic mutations in cell adhesion and chromatin remodeling genes. Nat Genet 2012, 44: 570–574.

8. Bashashati A, Haffari G, Ding J, Ha G, Lui K, Rosner J, Huntsman DG, Caldas C, Aparicio SA, Shah SP: DriverNet: uncovering the impact of somatic driver mutations on transcriptional networks in cancer. Genome Biol 2012, 13:R124.

9. Hou JP, Ma J: DawnRank: discovering personalized driver genes in cancer. Genome Med 2014, 6: 56.

10. Bertrand D, Chng KR, Sherbaf FG, Kiesel A, Chia BKH, Sia YY, Huang SK, Hoon DSB, Liu ET, Hillmer A, Nagarajan N: Patient-specific driver gene prediction and risk assessment through integrated network analysis of cancer omics profiles. Nucleic Acids Res 2015, 43: e44.

11. Vogelstein B, Papadopoulos N, Velculescu VE, Zhou S, Diaz LA, Kinzler KW: Cancer genome landscapes. Science 2013, 339: 1546–58.

12. Bhatia S, Frangioni J V, Hoffman RM, Iafrate AJ, Polyak K: The challenges posed by cancer heterogeneity. Nat Biotechnol 2012, 30: 604–10.

13. Lawrence MS, Stojanov P, Mermel CH, Robinson JT, Garraway LA, Golub TR, Meyerson M,Gabriel SB, Lander ES, Getz G: Discovery and saturation analysis of cancer genes across 21 tumour types. Nature 2014, 505: 495–501.

14. Garraway LA: Genomics-driven oncology: framework for an emerging paradigm. J Clin Oncol 2013, 31: 1806–14.

15. Soon WW, Hariharan M, Snyder MP: High-throughput sequencing for biology and medicine. Mol Syst Biol 2013, 9: 640.

16. Garay JP, Gray JW: Omics and therapy - a basis for precision medicine. Mol Oncol 2012, 6: 128–39.

17. Hortobagyi GN: Trastuzumab in the Treatment of Breast Cancer. N Engl J Med 2005, 353: 1734–1736.

18. Gunturu KS, Woo Y, Beaubier N, Remotti HE, Saif MW: Gastric cancer and trastuzumab: first biologic therapy in gastric cancer. Ther Adv Med Oncol2013, 5: 143–51.

19. Gonzalez-Perez A, Mustonen V, Reva B, Ritchie GRS, Creixell P, Karchin R, Vazquez M, Fink JL, Kassahn KS, Pearson J V, Bader GD, Boutros PC, Muthuswamy L, Ouellette BFF, Reimand J, Linding R, Shibata T, Valencia A, Butler A, Dronov S, Flicek P, Shannon NB, Carter H, Ding L, Sander C, Stuart JM, Stein LD, Lopez-Bigas N: Computational approaches to identify functional genetic variants in cancer genomes. Nat Methods 2013, 10: 723–9.

20. Kristensen VN, Lingjærde OC, Russnes HG, Vollan HKM, Frigessi A, Børresen-Dale A-L: Principles and methods of integrative genomic analyses in cancer. Nat Rev Cancer 2014, 14: 299–313.

21. Cheng F, Zhao J, Zhao Z: Advances in computational approaches for prioritizing driver mutations and significantly mutated genes in cancer genomes. Brief Bioinform 2015:bbv068–.

22. Ng PC: SIFT: predicting amino acid changes that affect protein function. Nucleic Acids Res 2003, 31: 3812–3814.

23. Adzhubei IA, Schmidt S, Peshkin L, Ramensky VE, Gerasimova A, Bork P, Kondrashov AS, Sunyaev SR: A method and server for predicting damaging missense mutations. Nat Methods 2010, 7: 248–9.

24. Schwarz JM, Cooper DN, Schuelke M, Seelow D: MutationTaster2: mutation prediction for the deep-sequencing age. Nat Methods 2014, 11: 361–362.

25. Reva B, Antipin Y, Sander C: Predicting the functional impact of protein mutations: application to cancer genomics. Nucleic Acids Res 2011, 39: e118.

26. Carter H, Chen S, Isik L, Tyekucheva S, Velculescu VE, Kinzler KW, Vogelstein B, Karchin R: Cancer-specific high-throughput annotation of somatic mutations: computational prediction of driver missense mutations. Cancer Res 2009, 69: 6660–7.

27. Gonzalez-Perez A, Deu-Pons J, Lopez-Bigas N: Improving the prediction of the functional impact of cancer mutations by baseline tolerance transformation. Genome Med 2012, 4: 89.

28. Shihab HA, Gough J, Cooper DN, Stenson PD, Barker GLA, Edwards KJ, Day INM, Gaunt TR: Predicting the functional, molecular, and phenotypic consequences of amino acid substitutions using hidden Markov models. Hum Mutat 2013, 34: 57–65.

29. Martelotto LG, Ng CK, De Filippo MR, Zhang Y, Piscuoglio S, Lim RS, Shen R, Norton L, Reis-Filho JS, Weigelt B: Benchmarking mutation effect prediction algorithms using functionally validated cancer-related missense mutations. Genome Biol 2014, 15: 484.

30. Gnad F, Baucom A, Mukhyala K, Manning G, Zhang Z: Assessment of computational methods for predicting the effects of missense mutations in human cancers. BMC Genomics 14 Suppl 3:S7.

31. Lawrence MS, Stojanov P, Polak P, Kryukov G V, Cibulskis K, Sivachenko A, Carter SL, Stewart C, Mermel CH, Roberts SA, Kiezun A, Hammerman PS, McKenna A, Drier Y, Zou L, Ramos AH, Pugh TJ, Stransky N, Helman E, Kim J, Sougnez C, Ambrogio L, Nickerson E, Shefler E, Cortés ML, Auclair D, Saksena G, Voet D, Noble M, DiCara D, et al.: Mutational heterogeneity in cancer and the search for new cancer-associated genes. Nature 2013, 499: 214–8.

32. Dees ND, Zhang Q, Kandoth C, Wendl MC, Schierding W, Koboldt DC, Mooney TB, Callaway MB, Dooling D, Mardis ER, Wilson RK, Ding L: MuSiC: identifying mutational significance in cancer genomes. Genome Res 2012, 22: 1589–98.

33. Tamborero D, Gonzalez-Perez A, Lopez-Bigas N: OncodriveCLUST: exploiting the positional clustering of somatic mutations to identify cancer genes. Bioinformatics 2013, 29: 2238–44.

34. Gonzalez-Perez A, Lopez-Bigas N: Functional impact bias reveals cancer drivers. Nucleic Acids Res 2012, 40: e169.

35. Mermel CH, Schumacher SE, Hill B, Meyerson ML, Beroukhim R, Getz G: GISTIC2.0 facilitates sensitive and confident localization of the targets of focal somatic copy-number alteration in human cancers. Genome Biol 2011, 12:R41.

36. Reimand J, Bader GD: Systematic analysis of somatic mutations in phosphorylation signaling predicts novel cancer drivers. Mol Syst Biol 2013, 9: 637.

37. Tokheim CJ, Papadopoulos N, Kinzler KW, Vogelstein B, Karchin R: Evaluating the evaluation of cancer driver genes. Proc Natl Acad Sci U S A 2016, 113: 14330–14335.

38. Akavia UD, Litvin O, Kim J, Sanchez-Garcia F, Kotliar D, Causton HC, Pochanard P, Mozes E, Garraway LA, Pe’er D: An integrated approach to uncover drivers of cancer. Cell 2010, 143: 1005–17.

39. Tamborero D, Lopez-Bigas N, Gonzalez-Perez A: Oncodrive-CIS: a method to reveal likely driver genes based on the impact of their copy number changes on expression. PLoS One 2013, 8: e55489.

40. Hautaniemi S, Ringnér M, Kauraniemi P, Autio R, Edgren H, Yli-Haija O, Astola J, Kallioniemi A, Kallioniemi O-P: A strategy for identifying putative causes of gene expression variation in human cancers. J Franklin Inst 2004, 341: 77–88.

41. Louhimo R, Lepikhova T, Monni O, Hautaniemi S: Comparative analysis of algorithms for integration of copy number and expression data. Nat Methods 2012, 9: 351–5.

42. Babur Ö, Gönen M, Aksoy BA, Schultz N, Ciriello G, Sander C, Demir E: Systematic identification of cancer driving signaling pathways based on mutual exclusivity of genomic alterations. Genome Biol 2015, 16: 45.

43. Leiserson MDM, Blokh D, Sharan R, Raphael BJ: Simultaneous identification of multiple driver pathways in cancer. PLoS Comput Biol 2013, 9: e1003054.

44. Cerami E, Demir E, Schultz N, Taylor BS, Sander C: Automated network analysis identifies core pathways in glioblastoma. PLoS One 2010, 5: e8918.

45. Leiserson MDM, Vandin F, Wu H-T, Dobson JR, Eldridge J V, Thomas JL, Papoutsaki A, Kim Y, Niu B, McLellan M, Lawrence MS, Gonzalez-Perez A, Tamborero D, Cheng Y, Ryslik GA, Lopez-Bigas N, Getz G, Ding L, Raphael BJ: Pan-cancer network analysis identifies combinations of rare somatic mutations across pathways and protein complexes. Nat Genet 2014, 47: 106–114.

46. Ding L, Wendl MC, McMichael JF, Raphael BJ: Expanding the computational toolbox for mining cancer genomes. Nat Rev Genet 2014, 15: 556–570.

47. Futreal PA, Coin L, Marshall M, Down T, Hubbard T, Wooster R, Rahman N, Stratton MR: A census of human cancer genes. Nat Rev Cancer 2004, 4: 177–83.

48. Santarius T, Shipley J, Brewer D, Stratton MR, Cooper CS: A census of amplified and overexpressed human cancer genes. Nat Rev Cancer 2010, 10: 59–64.

49. UniProt: a hub for protein information. Nucleic Acids Res 2015, 43:D204–D212.

50. Pletscher-Frankild S, Pallejà A, Tsafou K, Binder JX, Jensen LJ: DISEASES: Text mining and data integration of disease–gene associations. Methods 2015, 74: 83–89.

51. An O, Pendino V, D’Antonio M, Ratti E, Gentilini M, Ciccarelli FD: NCG 4.0: the network of cancer genes in the era of massive mutational screenings of cancer genomes. Database (Oxford) 2014, 2014: bau015.

52. Rubio-Perez C, Tamborero D, Schroeder MP, Antolín AA, Deu-Pons J, Perez-Llamas C, Mestres J, Gonzalez-Perez A, Lopez-Bigas N: In Silico Prescription of Anticancer Drugs to Cohorts of 28 Tumor Types Reveals Targeting Opportunities. Cancer Cell 2015, 27: 382–396.

53. Tamborero D, Gonzalez-Perez A, Perez-Llamas C, Deu-Pons J, Kandoth C, Reimand J, Lawrence MS, Getz G, Bader GD, Ding L, Lopez-Bigas N: Comprehensive identification of mutational cancer driver genes across 12 tumor types. Sci Rep 2013, 3: 2650.

54. Redig AJ, Jänne PA: Basket trials and the evolution of clinical trial design in an era of genomic medicine. J Clin Oncol 2015, 33: 975–7.

55. Miller DG: On the nature of susceptibility to cancer. The presidential address. Cancer 1980, 46: 1307–1318.

56. Schinzel AC: Oncogenic transformation and experimental models of human cancer. Front Biosci 2008, 13: 71.

57. OncoKB [http://oncokb.org/#/]

58. Ascierto PA, Kirkwood JM, Grob J-J, Simeone E, Grimaldi AM, Maio M, Palmieri G, Testori A, Marincola FM, Mozzillo N: The role of BRAF V600 mutation in melanoma. J Transl Med 2012, 10: 85.

59. Massacesi C, Di Tomaso E, Urban P, Germa C, Quadt C, Trandafir L, Aimone P, Fretault N, Dharan B, Tavorath R, Hirawat S: PI3K inhibitors as new cancer therapeutics: implications for clinical trial design. Onco Targets Ther 2016, 9: 203–10.

60. Dwork C, Kumar R, Naor M, Sivakumar D: Rank aggregation methods for the Web. In Proceedings of the tenth international conference on World Wide Web - WWW ’01. New York, New York, USA: ACM Press; 2001: 613–622.

61. Cheng W-C, Chung I-F, Chen C-Y, Sun H-J, Fen J-J, Tang W-C, Chang T-Y, Wong T-T, Wang H-W: DriverDB: an exome sequencing database for cancer driver gene identification. Nucleic Acids Res 2014, 42(Database issue):D1048–54.

62. Cowley GS, Weir BA, Vazquez F, Tamayo P, Scott JA, Rusin S, East-Seletsky A, Ali LD, Gerath WF, Pantel SE, Lizotte PH, Jiang G, Hsiao J, Tsherniak A, Dwinell E, Aoyama S, Okamoto M, Harrington W, Gelfand E, Green TM, Tomko MJ, Gopal S, Wong TC, Li H, Howell S, Stransky N, Liefeld T, Jang D, Bistline J, Hill Meyers B, et al.: Parallel genome-scale loss of function screens in 216 cancer cell lines for the identification of context-specific genetic dependencies. Sci Data 1: 140035.

63. Barretina J, Caponigro G, Stransky N, Venkatesan K, Margolin AA, Kim S, Wilson CJ, Lehár J, Kryukov G V, Sonkin D, Reddy A, Liu M, Murray L, Berger MF, Monahan JE, Morais P, Meltzer J, Korejwa A, Jané-Valbuena J, Mapa FA, Thibault J, Bric-Furlong E, Raman P, Shipway A, Engels IH, Cheng J, Yu GK, Yu J, Aspesi P, de Silva M, et al.: The Cancer Cell Line Encyclopedia enables predictive modelling of anticancer drug sensitivity. Nature 2012, 483: 603–7.

64. Garnett MJ, Edelman EJ, Heidorn SJ, Greenman CD, Dastur A, Lau KW, Greninger P, Thompson IR, Luo X, Soares J, Liu Q, Iorio F, Surdez D, Chen L, Milano RJ, Bignell GR, Tam AT, Davies H, Stevenson JA, Barthorpe S, Lutz SR, Kogera F, Lawrence K, McLaren-Douglas A, Mitropoulos X, Mironenko T, Thi H, Richardson L, Zhou W, Jewitt F, et al.: Systematic identification of genomic markers of drug sensitivity in cancer cells. Nature 2012, 483: 570–5.

65. Zhao J, Zhang S, Wu L-Y, Zhang X-S: Efficient methods for identifying mutated driver pathways in cancer. Bioinformatics 2012, 28: 2940–2947.

66. Youn A, Simon R: Identifying cancer driver genes in tumor genome sequencing studies. Bioinformatics 2011, 27: 175–181.

67. Ciriello G, Cerami E, Sander C, Schultz N: Mutual exclusivity analysis identifies oncogenic network modules. Genome Res 2012, 22: 398–406.

68. Ng S, Collisson EA, Sokolov A, Goldstein T, Gonzalez-Perez A, Lopez-Bigas N, Benz C, Haussler D, Stuart JM: PARADIGM-SHIFT predicts the function of mutations in multiple cancers using pathway impact analysis. Bioinformatics 2012, 28:i640–i646.

